# Genome-Scale Codon Deoptimization Enables Attenuation of Rift Valley Fever Virus

**DOI:** 10.64898/2026.08.13.744589

**Authors:** Sandra Moreno, Alejandro Cenalmor, Celia Alonso, Gema Lorenzo, Carlos J. Ciria-Gil, Belén Borrego, Luis Martinez-Sobrido, Alejandro Brun, Aitor Nogales

**Author notes:** Contributed equally (Sandra Moreno, Alejandro Cenalmor, Celia Alonso).

## Abstract

Rift Valley Fever Virus (RVFV) is a mosquito-borne zoonotic pathogen responsible for severe disease in domestic and wild ungulates as well as humans, representing a major threat to livestock production and human public health. RVFV is endemic in many African countries and has the potential to spread to new geographical regions. Current vaccines have limitations in safety and efficacy, highlighting the need for strategies to develop new vaccines candidates. In this study, we explored the use of codon deoptimization (CD) as a novel attenuation approach for the development of live-attenuated vaccine (LAV) against RVFV. CD exploits the redundancy of the genetic code by replacing frequently used codons with synonymous, less-preferred codons, thereby reducing translational efficiency without altering the amino acid sequence. We recoded parts of the M and S genome segments of RVFV using the least frequently used codons in mammalian cells, ensuring complete preservation of protein functionality and immunogenicity. Using reverse genetics, we rescued a panel of recombinant (r)RVFV encoding codon-deoptimized S-segment NSs gene (rNScd), M-segment Gn/Gc genes (rMcd), or both (rMcd/NScd). These recombinant CD viruses were characterized *in vitro* in mammalian and insect cell lines and *in vivo* using wild-type and immunocompromised mice. Results demonstrated varying degrees of attenuation among the three CD rRVFV, with the one deoptimized in both viral segments, rMcd/NScd, as a promising LAV based on the safety profiles. This study provides proof of concept for the use of CD as a rational strategy to generate attenuated RVFV, for the development of next-generation vaccines against this zoonotic threat.

## INTRODUCTION

Rift Valley fever (RVF) is a re-emerging zoonotic disease endemic to Africa, the Middle East, and parts of the Indian Ocean region, causing recurrent outbreaks in domestic and wild ruminants with spillover into humans [1]. The causative agent, Rift Valley fever virus (RVFV), is a mosquito-borne phlebovirus (family *Phenuiviridae*) transmitted primarily by *Aedes* and *Culex* mosquito species, but also through direct contact with infected animal fluids [2,3]. While most human infections are mild or asymptomatic, severe cases can lead to hemorrhagic fever, or death. In livestock, RVFV induces high abortion rates and neonatal mortality, resulting in significant economic losses [1,2,4,5]. RVFV is classified as a potential biothreat and is listed as a select agent in the United States and as a priority pathogen by the World Health Organization (WHO) due to its epidemic potential and lack of effective vaccines or antivirals (https://www.who.int/activities/prioritizing-diseases-for-research-and-development-in-emergency-contexts). The virus has a tri-segmented, negative-sense, single-stranded RNA genome of different sizes: Large (L), Medium (M) and Small (S). The L-segment encodes the RNA-dependent RNA polymerase (RdRp, or L protein); the M-segment encodes the viral glycoproteins Gn and Gc and the nonstructural protein (NSm) as well as a Nsm/Gn fusion protein, often known as the 78 kDa protein; and the S-segment encodes the viral nucleoprotein (N) and the nonstructural NSs protein, a major virulence factor [6–9].

The mammalian genetic code is degenerate, meaning most amino acids are encoded by multiple synonymous codons. These codons are not used with equal frequency, either within a single genome or across different DNA/RNA genomes [10,11]. This variation, known as codon usage bias, reflects differences in how organisms preferentially use certain codons to incorporate the same amino acid into proteins [12–14]. Leveraging this phenomenon, researchers have developed strategies such as codon optimization (CO) and codon deoptimization (CD) to respectively enhance or reduce protein expression in various systems. Importantly, CD modifies only the nucleotide sequence without altering the amino acid composition, protein function, or immunogenic properties [15]. CD has been used in several RNA viruses, including influenza A virus [16,17], arenaviruses [18–20], respiratory syncytial virus (RSV) [21]; and DNA viruses like vaccinia virus (VACV) [22].

In this study, we reengineered the RVFV M and S genome segments by introducing the least-used human synonymous codons through *de novo* gene synthesis, ensuring complete preservation of the wild-type (WT) amino acid sequence and, consequently, the immunogenicity and functionality of the encoded viral proteins. Using reverse genetics, we generated three recombinant viruses carrying CD mutations in the M (rMcd), S (rNScd), or both (rMcd/NScd) viral segments. These recombinant viruses were comprehensively characterized *in vitro* in mammalian and insect cell lines and *in vivo* in two murine models of infection. Our results demonstrate that the rMcd/NScd is attenuated, suggesting the feasibility of implementing a CD-based approach, alone or in combinantion with other attenuation approaches, for the rational development of LAV for the prevention of RVFV infection.

## MATERIAL AND METHODS

### Cell lines

Vero E6, HEK293T, and A549 cells were cultured in Dulbecco’s modified Eagle medium (DMEM) supplemented with 5% fetal bovine serum (FBS), 1% nonessential amino acids, penicillin (100 U/mL), streptomycin (100 µg/mL), and 2 mM L-glutamine. Cultures were maintained at 37°C in a humidified atmosphere containing 5% CO₂. BSR-T7 cells, which constitutively express the bacteriophage T7 RNA polymerase [23,24], were grown under identical conditions, with the addition of Geneticin (1 mg/mL). C6/36 cells, derived from *Aedes albopictus,* were propagated in Eagle’s Minimum Essential Medium (MEM) supplemented with 10% FBS, 2 mM L-glutamine, gentamicin (50 µg/mL), and MEM vitamin solution (Sigma), and maintained at 28°C.

### Plasmids

All pBlueScript II SK (+) based plasmids to rescue a recombinant WT (rWT) RVFV (South African strain 56/74) were synthesized *de novo* by Biomatik and previously described [24,25]. Plasmids were designed to contain the viral cDNAs flanked upstream by a T7 RNA polymerase promoter sequence and downstream by the hepatitis delta virus ribozyme (HDVR) and T7 RNA polymerase terminator sequences [24]. To generate the CD viruses, a plasmid containing the CD NSs and WT N gene in segment S was synthesized *de novo* using the strategy described above. For segment M, the CD design targeted a region spanning nucleotides 429 to 3614, which includes Gn and Gc glycoprotein-coding sequences. This design was carefully optimized to avoid disrupting the translation of other viral proteins encoded within segment M, such as NSm and the 78 kDa NSm/Gn fusion protein.

### Recovery and characterization of recombinant viruses

Recombinant WT and CD RVFV were rescued in BSR-T7 cells and subsequently amplified in Vero E6 cells as previously described [24]. Viral stocks were generated and titrated by plaque assay under high-containment conditions. Replication kinetics were evaluated in Vero E6 and C6/36 cells infected at a multiplicity of infection (MOI) of 0.01, with viral titers determined by plaque assay at different time points [24–27]. Plaque morphology at 37°C was assessed by plaque assay following immunostaining as previously described [24], using an anti-RVFV rabbit pAb generated in the laboratory [27].

### Protein gel electrophoresis and Western blot analysis

Cell extracts from either mock- or virus-infected (MOI, 0.01) Vero E6 cells were lysed at 24 or 48 h p.i. in radioimmunoprecipitation assay (RIPA) buffer. Proteins were separated by denaturing SDS-PAGE electrophoresis and transferred to nitrocellulose membranes. Western blot analysis were performed with specific primary rabbit anti-RVFV pAb [27]. A mAb against actin (Sigma) was used as an internal loading control. Bound primary antibodies were detected with horseradish peroxidase (HRP)-conjugated secondary antibodies. Inmunoblots were quantified by densitometry using the ImageJ software. Protein bands were normalized to the level of actin expression.

### Molecular and functional analyses

Upon extraction of viral RNA from infected cells, the identity/stability of recombinant viruses was confirmed by RT-PCR using specific primers (**Table 1**) and restriction enzyme analysis. To assess host shutoff activity, HEK293T cells were co-transfected with plasmids expressing WT or CD NSs proteins together with a Gaussia luciferase reporter construct. Luciferase activity was measured 24 h post-transfection as an indicator of host gene expression inhibition.

**Table 1.**
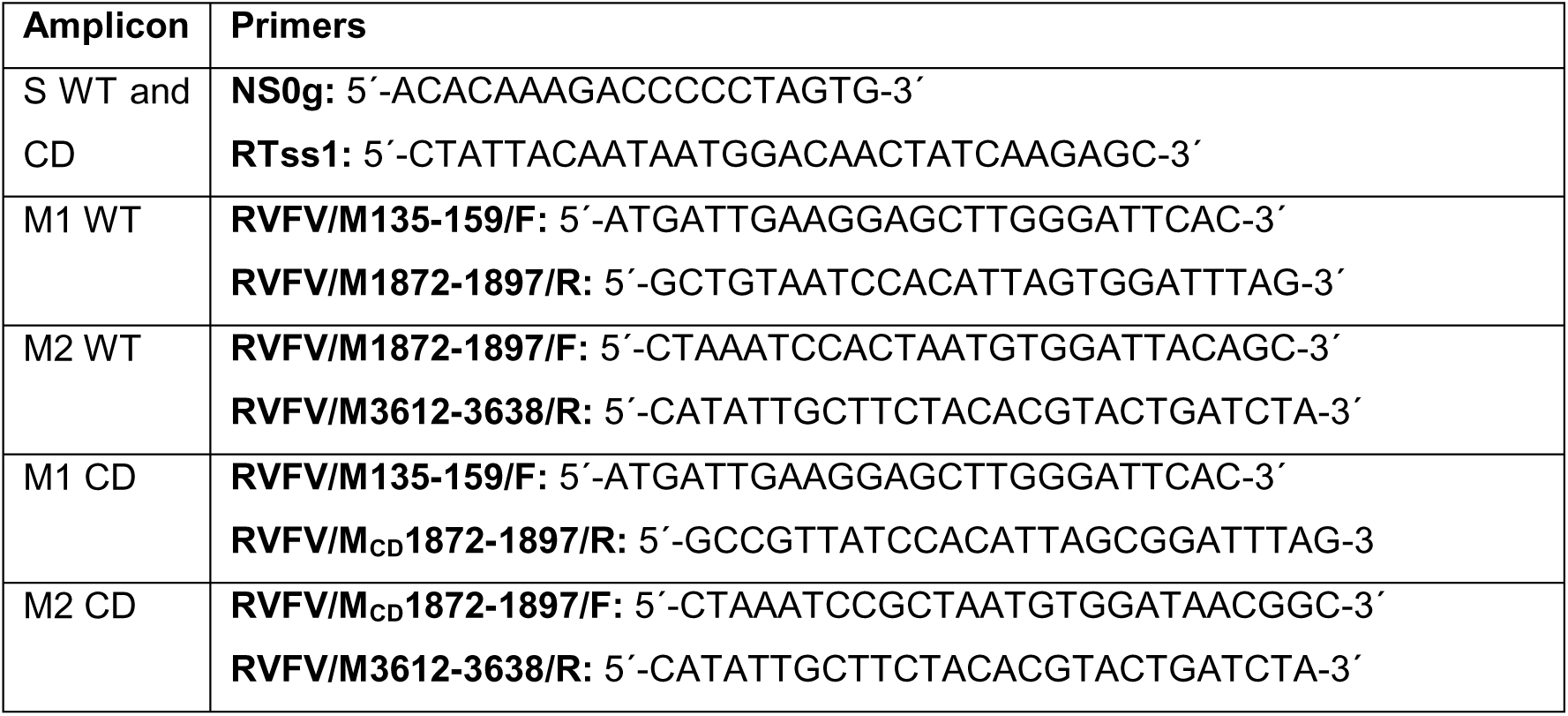
Primers used to amplified WT and CD segments.

### Animal studies and immunological assays

Pathogenicity was evaluated in 8- to 10-week-old 129/Sv (WT) and type-I interferon receptor-deficient A129 (IFNAR^⁻/⁻^) male mice (n = 5/group). Mice were housed and used at the biosafety level 3 (BSL-3) animal facility. Ethical approval was granted by Comunidad de Madrid veterinary authorities (PROEX 079.6/22). Animals were inoculated intraperitoneally with 10^4^ PFU of recombinant viruses and monitored for 14 days for clinical signs (such as malaise, respiratory distress, and lack of movement), body weight changes, and survival. Viremia was assessed by RT-qPCR [28] using blood samples collected by submandibular puncture at 72 h.p.i. Sera obtained at 15 days post-infection (d.p.i.) were analyzed by an in-house ELISA to detect anti-RVFV N antibodies, as described previously [24,26].

### Genome sequencing and statistical analysis

Viral genomes were characterized using sequence-independent single-primer amplification (SISPA) method described by Djikeng et al., [29], followed by Oxford Nanopore sequencing. Sequence assembly and analysis were performed using GENPAT, Geneious Prime, and IGV software. Statistical analyses were performed using GraphPad Prism and Microsoft Excel, with p-values ≤ 0.05 considered statistically significant.

## RESULTS

### Generation of CD recombinant RVFV

Codon-deoptimized versions of the RVFV M and NSs genes were designed using underrepresented codons based on the *Homo sapiens* codon usage, while preserving the original amino acid sequences (**Figure 1**). To do so we used the CoDe web-based bioinformatic tool (Sharma et al 2023). The M-segment encodes a single open reading frame (ORF), yet its mRNA can produce at least four distinct proteins through leaky scanning of five alternative initiation codons and co-translational cleavage of polypeptides [30,31]. To generate the CD M segment (Mcd), we introduced synonymous codon changes downstream of the fifth AUG initiation codon to CD Gc/Gn, avoiding disrupt the expression of other viral proteins encoded in segment M (**Figure 1A**).

**Figure 1.**
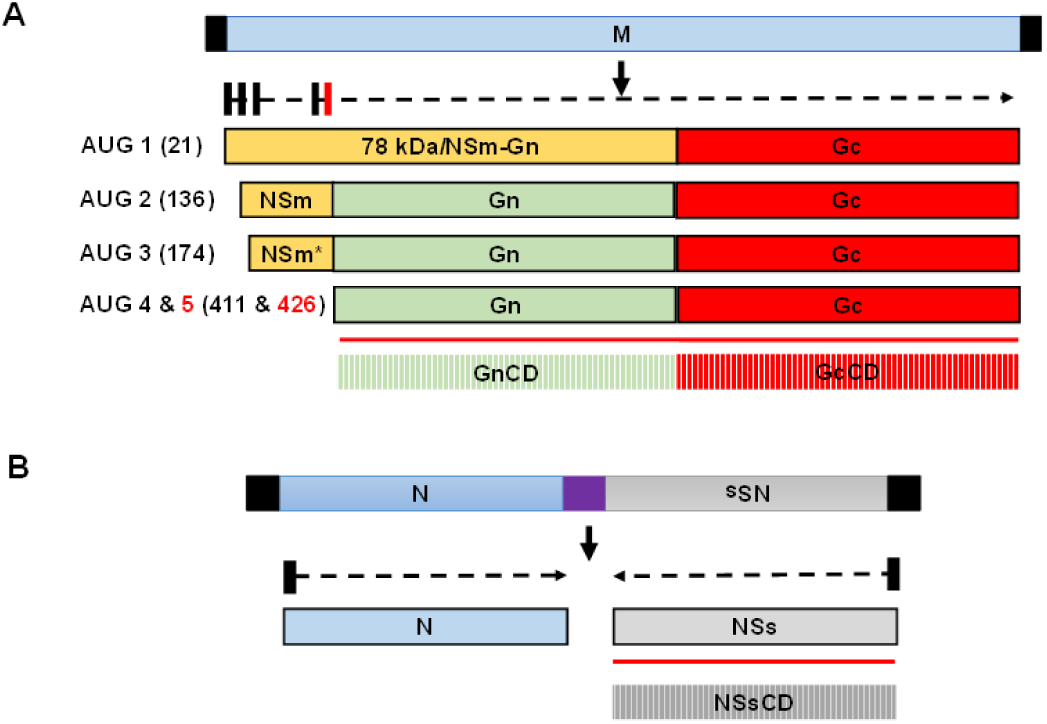
Schematic representation of the WT and CD RVFV M and S segments. (**A**) The relative nucleotide position of the different in-frame AUG codons in the NSm region and the coding regions of NSm, Gn and Gc are depicted on the upper graph with black or red boxes. This panel also shows the schematic representation of the polypeptides expressed from each of the AUG codons. Nucleotide positions are indicated in parentheses. Different proteins, including P78 and NSm and NSm* (orange), Gn (green) and Gc (red), are generated upon cleavage of polyprotein precursors. Untranslated regions (UTR) flank each segment (black boxes). The codon deoptimized region in this segment starts at the AUG 5 and highlighted by a red box in the dashed arrow. The red line indicates the deoptimized region of the protein (striped boxes). (**B**) In the S segment, the nucleoprotein (N) is generated by an mRNA transcribed from the negative-sense genomic viral RNA (vRNA), and the non-structural S protein (NSs) is produced from an mRNA transcribed from the antigenomic copy of vRNA generated during viral replication. Only the NSs protein (ORF) is codon deoptimized in this segment (NSs^CD^) and the codon deoptimization is represented with a red line. The purple box represents an intergenic transcription termination site in the S segment.

A total of 69.6% of codons in the Gn/Gc genes and 73.5% in the NSs gene were modified, resulting in reduced adaptation to mammalian codon usage without major changes in GC content. We also calculated the codon adaptation index (CAI) for WT and CD sequences using codon usage tables for *Homo sapiens* and *Drosophila melanogaster* (**Table 2a**). Interestingly, WT sequences were more adapted to human hosts than to insect hosts for both genes, whereas CD significantly reduced the CAI for human codon usage. In contrast, CD had little effect on CAI values for insect codon usage (**Table 2a**). Of note, the percentages of CpG (and UpA) dinucleotides increased with respect to the WT sequences (**Table 2b**).

**Table 2a.** Synonymous nucleotide or codon substitutions and CAI calculation.

| Viral gene | Nucleotide |  |  | Codon |  | CAI index |  |  |  |
| --- | --- | --- | --- | --- | --- | --- | --- | --- | --- |
|  | Mutations (total) | % | GC % (WT/CD) | Mutations | % | WT (Hs) | CD (Hs) | WT (Dm) | CD (Dm) |
| **M** | 865 (3480) | 24.9 | 46.3/46.4 | 807 (1160) | 69.6 | 0.74 | 0.45 | 0.57 | 0.51 |
| **NSs** | 225 (798) | 28.2 | 47.2/<br>44.6 | 191 (260) | 73.5 | 0.74 | 0.43 | 0.61 | 0.52 |

CAI index: (https://www.biologicscorp.com/tools/CAICalculator). Hs (Homo Sapiens), Dm (Drosophila melanogaster).

**Table 2b.** Changes in the frequency of CpG and UpA dinucleotides.

| segment | GC% | Total CpG<br>(difference)* | Total UpA<br>(difference)* | Ratio†<br>CpG | Ratio†<br>UpA |
| --- | --- | --- | --- | --- | --- |
| M-WT | 46.3 | 39 | 184 | 0.19 | 0.63 |
| M-CD | 46.4 | 333 (+294) | 366 (+182) | 1.95 | 1.16 |
| NSs-WT | 47.2 | 35 | 69 | 0.34 | 0.63 |
| NSs-CD | 44.6 | 103 (+68) | 118 (+49) | 1.05 | 1.03 |

\* Additional dinucleotides in the mutated sequence with respect to WT.
† Observed-to-expected ratio of dinucleotide frequency, corrected for GC content.

Recombinant viruses carrying CD mutations in M (rMcd), NSs (rNScd), or both genes (rMcd/NScd) were successfully rescued using our previously described reverse genetics approach [24]. All viruses were plaque-purified and viral stocks generated from two independent plaques for each CD virus. Identity and integrity of the engineered genomic segments were confirmed by PCR and restriction enzyme analysis using CD-specific markers (**Figure 2**).

**Figure 2.**
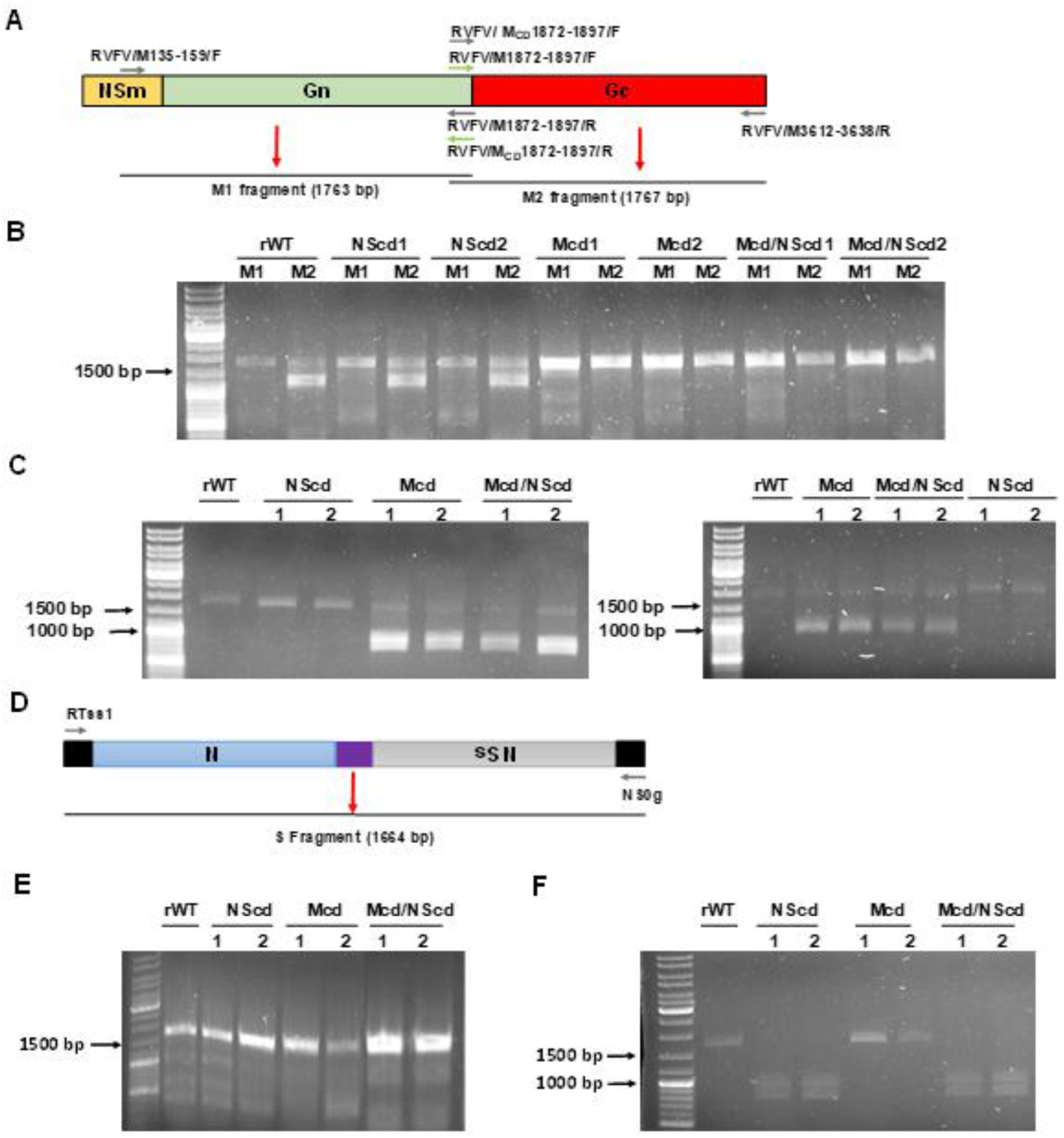
Genome characterization of M and S segments. Vero E6 cells were infected (MOI of 0.01) with rWT or with the CD viruses expressing NS_CD_, M_CD_, and M/NS_CD_, and total RNA was extracted. (**A and B**) The RVFV M segment was obtained by PCR using specific WT (grey arrows) or CD (green arrows) primers designed to amplify two regions (M1 and M2) of the segment M. An unspecific or double band was observed only for M2 when amplified with primers specific for the WT sequence, further confirming the identity of the sequence. (**C**) Restriction pattern assay. Fragments M1 and M2 (WT and CD) were digested using the restriction enzymes XbaI for M1 (953 + 810 bp) and EcoRI for M2 (851+916 bp), which cut only the CD fragments. (**D and E**) The S segment was analyzed using specific primers (common for WT and CD sequences). The S segment was digested using the restriction enzyme KpnI (914+750 bp), which cut only the CD sequence (**F**).

Segment M was amplified in two overlapping fragments (positions 135– 1897 and 1872–3638) using WT- or CD-specific primers. As expected, amplification occurred only when primers matched the respective WT or CD sequence (**Figures 2A and B**). Fragments 1 and 2 were then digested with XbaI and EcoRI restriction enzymes, respectively, since these restriction sites were engineered into the CD sequence. The digestion patterns confirmed the identity of CD viruses, as only CD fragments were cleaved (**Figure 2C**). Similarly, the entire S-segment was amplified using specific primers (**Figure 2D**), and digested with KpnI restriction enzyme, which recognizes a site introduced in the CD NSs sequence (**Figures 2E and F**). The restriction patterns validated the identity of the recombinant viruses containing the CD NSs gene.

### *In vitro* characterization of CD recombinant RVFV in mammalian cells

Replication kinetics in Vero E6 cells (**Figures 3A-3C**) revealed that rNScd viruses (rNScd-1 and rNScd-2) replicated similarly to rWT RVFV, indicating that CD of NSs alone did not significantly affect viral fitness (**Figure 3A**). In contrast, viruses carrying the Mcd segment (rMcd-1 and rMcd-2) (**Figure 3B)**, either alone or combined with NScd (Mcd/NScd-1 and Mcd/NScd-2) (**Figure 3C**), showed reduced growth and lower peak viral titers. Consistent with these findings, Mcd-containing viruses produced smaller plaques than rWT and rNScd viruses in Vero E6 cells (**Figure 3D**). The RVFV NSs protein, a key virulence factor, suppresses the host innate antiviral response to facilitate viral replication and dissemination [32–34]. To evaluate whether CD of NSs affects this function, mammalian IFN-competent A549 cells were infected (MOI 0.01) (**Figure 3E**). Infection of A549 cells yielded similar results, confirming that attenuation associated with Mcd was independent of the host antiviral response.

**Figure 3.**
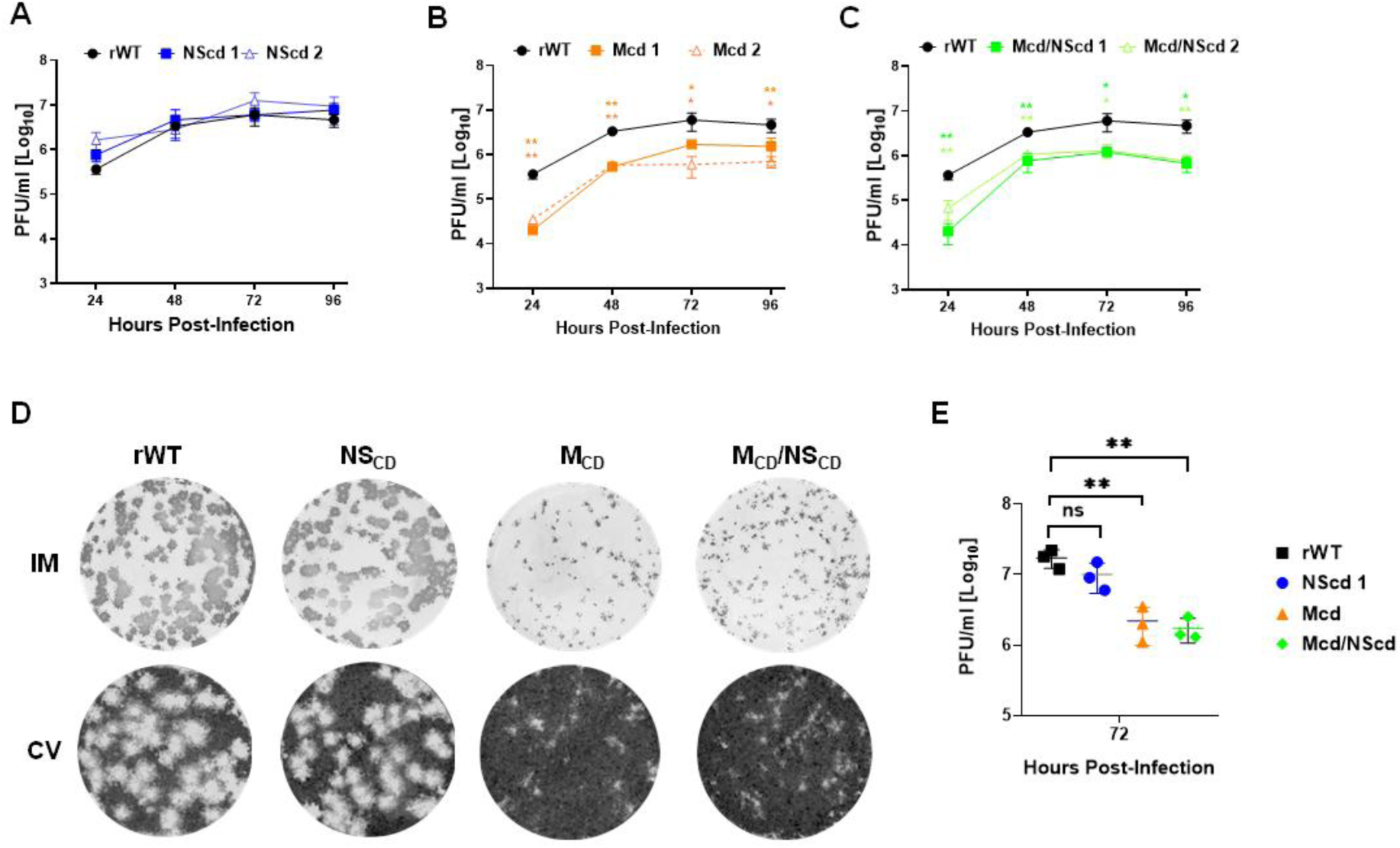
Characterization of recombinant RVFVs in cell culture. (A, B and C) Multicycle growth kinetics. Vero E6 cells were infected (MOI of 0.01) with rWT or two clones for each CD virus: rNScd, rMcd and rMcd/NScd. At the indicated h p.i., viral titers in culture supernatants were evaluated by plaque assay (PFU/mL). The same rWT representation was used in A, B and C. **(D) Plaque assays.** Vero E6 cells were infected with rWT or one selected clone for each CD virus: rNS_CD_, rM_CD_ and rM_CD_/NS_CD_. and incubated at 37°C for 3 days. Plaques were evaluated by immunostaining **(top)** and then, crystal violet stained **(bottom). (E) Replication in A549 cells.** A549 cells were infected (MOI of 0.01) with rWT or rCD virus. Then, at 72 h p.i., viral titers in culture supernatants were evaluated by plaque assay (PFU/mL). Data (A, B, C, and D) represent the means ± SDs of triplicate samples. The differences between the groups were calculated using one-way ANOVA with the Dunnett’s multiple comparisons test, *P < 0.05; **P < 0.01.

### Analysis of protein expression in cells infected with CD recombinant RVFV

Western blot analysis of infected Vero E6 cells demonstrated reduced expression of the viral glycoproteins Gn and Gc in rMcd and rrMcd/NScd viruses, particularly at 24 h post-infection, while N expression remained unaffected (**Figure 4**). Western blot analysis revealed distinct bands corresponding to the expected molecular sizes of Gn (≈54 kDa), Gc (≈56 kDa), and N (≈25 kDa) proteins. These results suggest that reduced glycoprotein expression contributes to the impaired replication of Mcd-containing viruses. Notably, no differences were observed between rWT and rNScd viruses (**Figure 4**).

**Figure 4.**
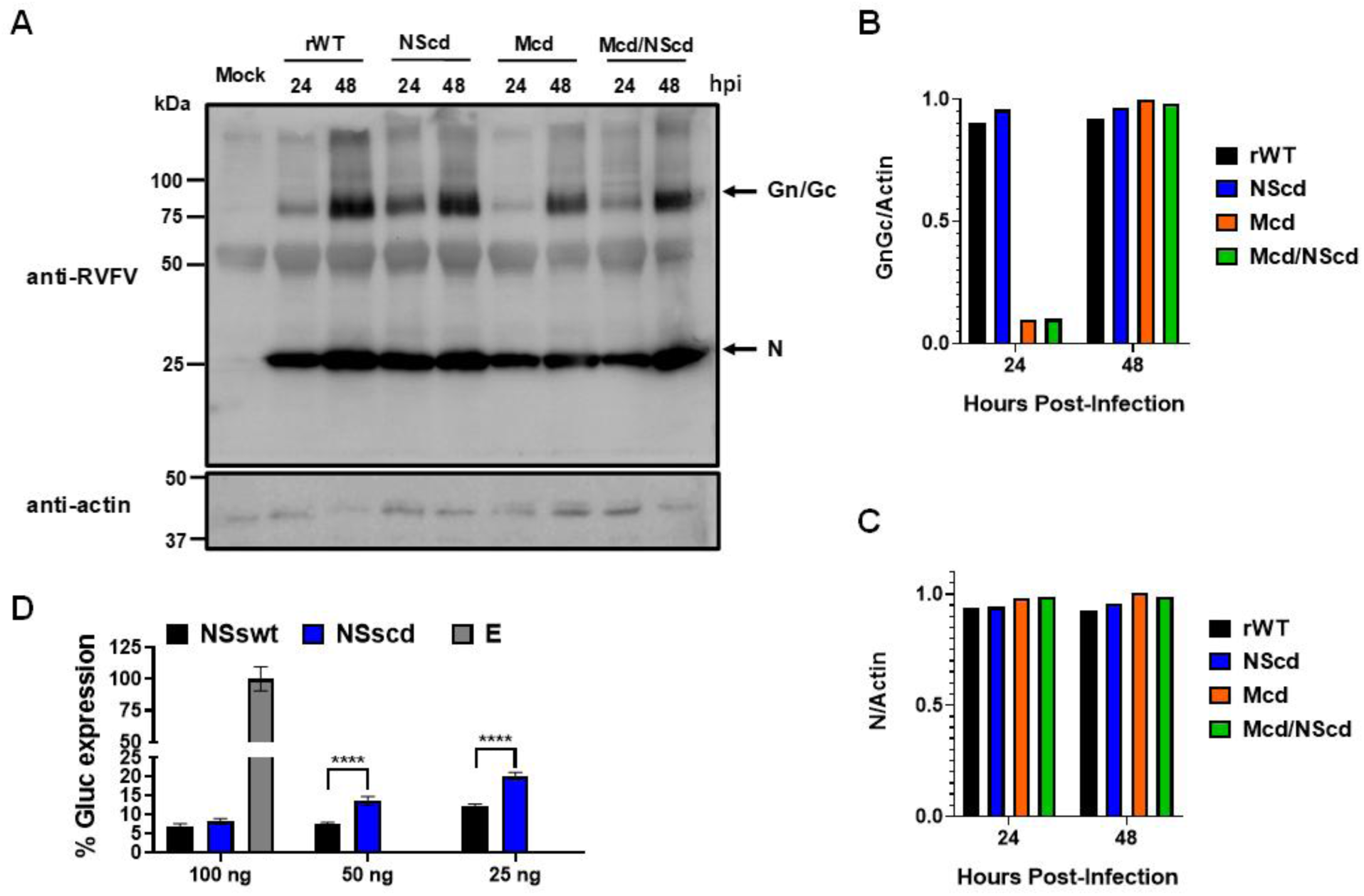
Analysis of protein expression by WB. **(A)** Vero E6 cells were mock-infected or infected (MOI of 0.01) with rWT or each CD virus: rNScd, rMcd and rMcd/NScd. At 24 or 48 h p.i, protein expression was examined by Western blotting using specific antibodies for viral proteins: pAb against RVFV for N and Gn. Actin was used as a loading control. The numbers on the left indicate the molecular size of the protein markers (in kilodaltons, kDa). (**B and C**) Western blots were quantified by densitometry. Protein bands were normalized to the level of actin expression. (**D**) Inhibition of host gene expression by NSs. Human HEK293Twere cotransfected with 0, 25, 50, or 100 ng of pCAGGS expression plasmids encoding NSs fused to an HA epitope tag, together with 15 ng of pCAGGS plasmid encoding Gluc. Empty plasmid was included as a control. At 24 h pt, Gluc activity was quantified using a luciferase plate reader. Statistical significance was calculated using one-way ANOVA (*P < 0.05; **P < 0.01; ****P < 0.0001)".

To evaluate the expression of NSs we performed a functional assay to determine whether the codon-deoptimized NSs protein is able to inhibit host gene expression, as described before [32–35]. Functional analysis of NSs-mediated host shutoff activity showed that both WT and codon-deoptimized NSs inhibited cellular gene expression in a dose-dependent manner. Empty plasmid was included as a negative control. However, the CD NSs construct displayed reduced inhibitory activity, consistent with lower protein expression levels (**Figure 4D**).

### Growth properties of CD recombinant RVFV in C6/36 mosquito cells

To assess whether CD influences viral replication in insect cells, recombinant viruses were evaluated in C6/36 mosquito cells infected (MOI 0.01) with rWT, rMcd, rNScd, or rMcd/NScd, and viral titers were quantified (**Figure 5**). As observed in mammalian cells (**Figure 3**), viruses containing the Mcd segment replicated less efficiently than rWT and rNScd viruses, whereas codon deoptimization of NSs alone had little effect. These results indicate that attenuation associated with the M segment is maintained across both mammalian and insect hosts. This can be explained based on the differences in CAI values for WT and CD M segments, between human and insects (**Table 2a**).

**Figure 5.**
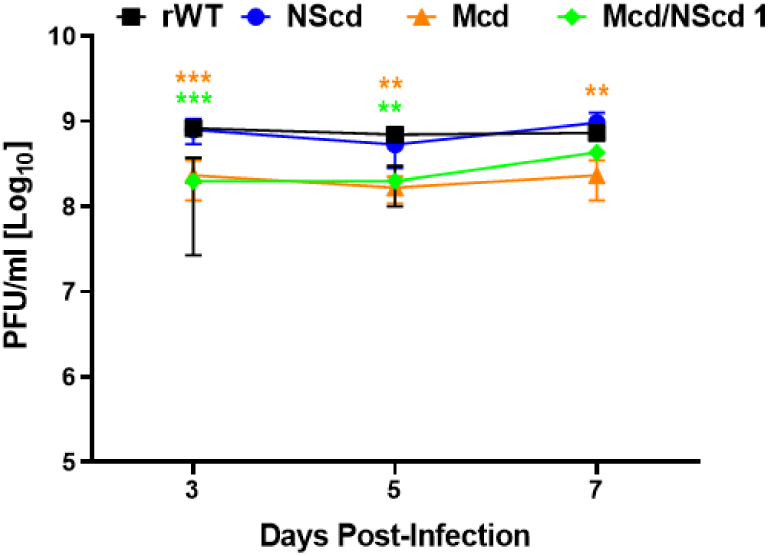
Multicycle growth kinetics in insect cells. C6/36 mosquito cells were infected (MOI of 0.01) with rWT or CD virus: rNScd, rMcd and rMcd/NScd. At the indicated 3, 5 and 7 days p.i., viral titers in culture supernatants were evaluated by plaque assay (PFU/mL). Data represent the means ± SDs of triplicate samples. The differences between the groups were calculated using two-way ANOVA with the Dunnett’s multiple comparisons test, ***P < 0.001; **P < 0.01.

### Pathogenicity and replication of CD recombinant RVFV in mice

The virulence of recombinant viruses was assessed in WT 129/Sv mice (**Figure 6**) and highly susceptible IFNAR^⁻/⁻^ A129 mice (**Figure 7**). In WT animals, all CD viruses showed varying degrees of attenuation compared with rWT. Mice infected with rWT succumbed rapidly, whereas rNScd-infected animals survived substantially longer. The strongest attenuation was observed for rMcd/NScd, with only one animal succumbing to infection. Clinical signs of infection (**Figure 6C**) were also recorded daily and correlated with body weight and survival data. Viremia at 3 d p.i. (**Figure 6D**) was significantly higher in rMcd-infected animals, as expected since rWT-infected mice died rapidly and rNScd and rMcd/NScd were significantly attenuated as compared with rWT or rMcd. Notably, all surviving animals developed RVFV-specific anti-N antibodies (**Figure 6E**), a standard marker of RVFV infection and replication [26,27,36–38], and neutralizing responses with protective levels [32] (data not shown), confirming productive infection and seroconversion. Notably, although rNScd exhibited little attenuation in cell culture, significant attenuation was evident in vivo, suggesting that codon deoptimization of NSs influences virulence through mechanisms not captured in vitro.

**Figure 6.**
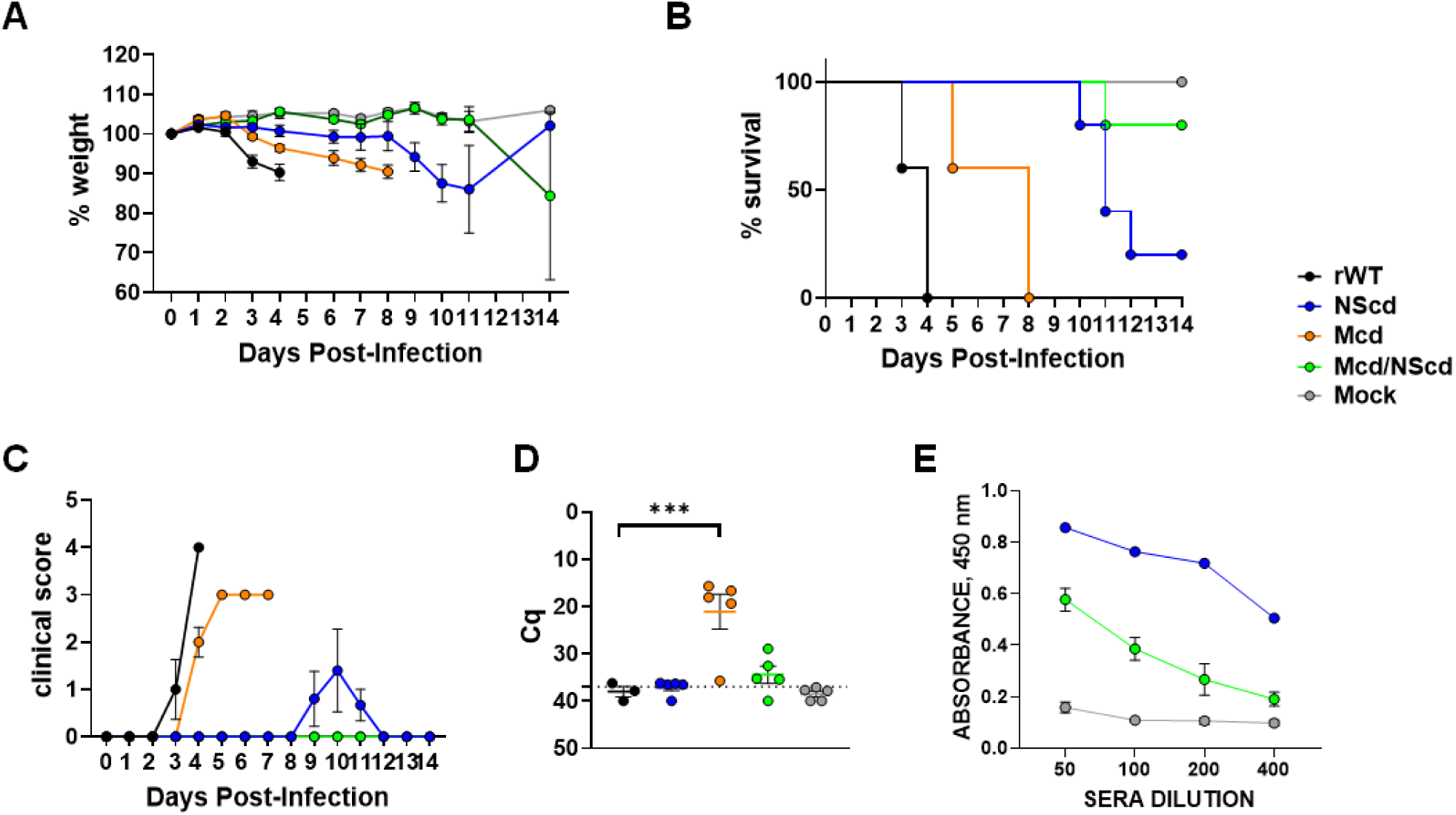
*In vivo* virulence of rRVFV in WT mice. Groups of 8- to 12-week-old WT A129 male mice (N = 5/group) were infected with the indicated viruses: rWT or the CD viruses: rNScd, rMcd and rMcd/NScd. Weight loss (**A**) and survival (**B**) were evaluated daily for 14 days. (**C**) Clinical score represented as a cumulative total assigned for each sign recorded. (**D**) Viremia was determined in blood samples collected at day 3 p.i. Cutoff: Cq ≥37 (dotted gray line). Points represent individual Cq values for each mouse, and lines of the corresponding color represent the mean Cq values of each group. Dunnett’s method was used to statistically compare CD viruses versus rWT, \*\*\**P* < 0.0002. (**E**) Antibody responses in surviving mice were evaluated by N-ELISA on day 15 p.i., to confirm seroconversion.

**Figure 7.**
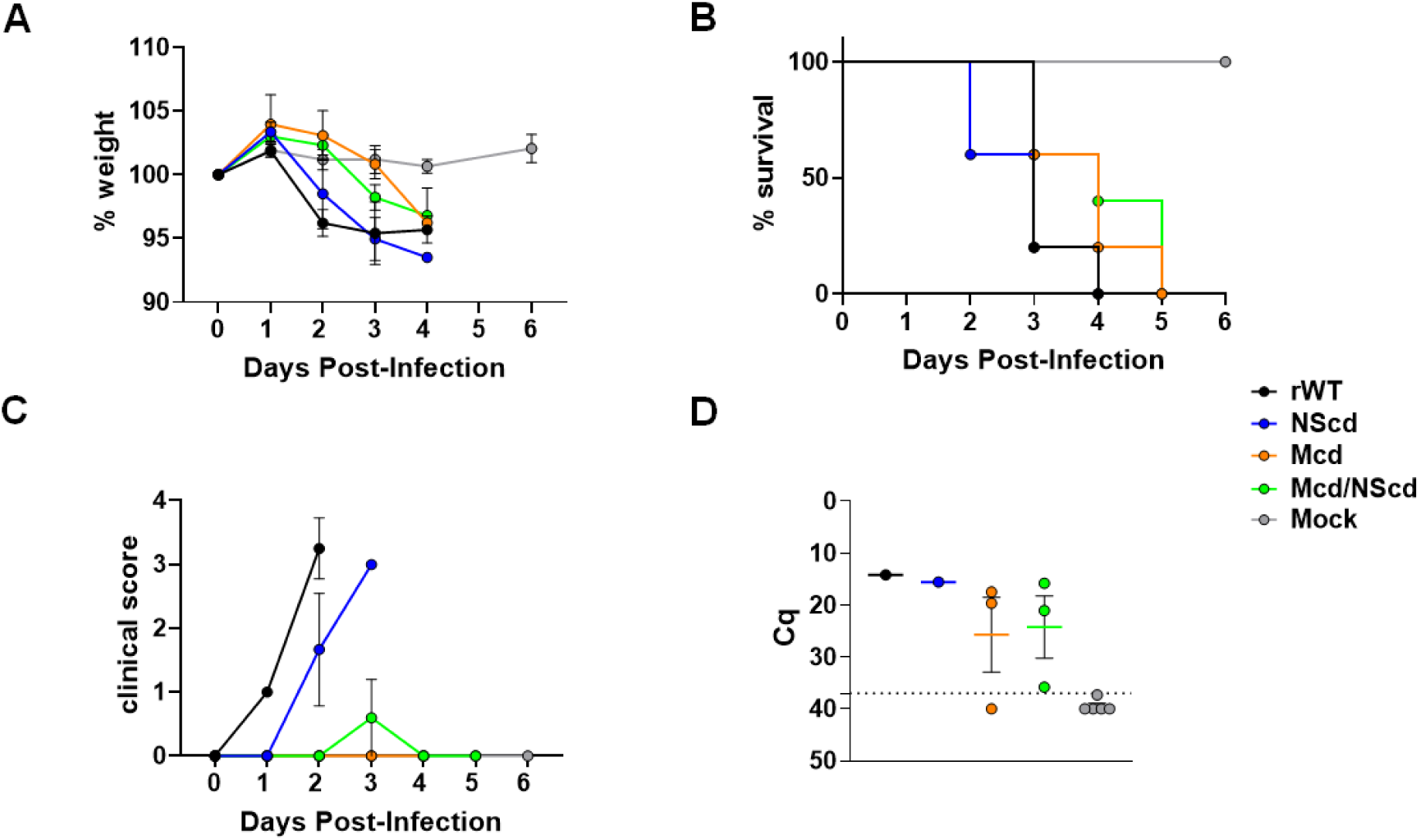
*In vivo* virulence of rRVFV in A129 IFNAR^-/-^ mice. Groups of 8- to 12-week-old IFNAR-/- A129 male mice (N = 5/group) were infected with the indicated viruses: rWT or the CD viruses: rNScd, rMcd and rMcd/NScd. Weight loss (**A**) and survival (**B**) were evaluated daily for 6 days. (**C**) Clinical score represented as a cumulative total assigned for each sign recorded. (**D**) Viremia was determined in blood samples collected at day 3 p.i. Cutoff: Cq ≥37 (dotted gray line). Points represent individual Cq values for each mouse, and lines of the corresponding color represent the mean Cq values of each group. Dunnett’s method was used to statistically compare rCD viruses versus rWT and non-significant differences were observed.

In IFNAR^−/−^ mice (**Figure 7**), which are highly susceptible to RVFV infection, all viruses remained lethal, although animals infected with rMcd or rMcd/NScd exhibited reduced clinical severity compared with those infected with rWT.

### Genome Stability Analysis

Nanopore sequencing of viral stocks of two independent clones for each virus revealed that CD viruses accumulated more mutations than rWT, predominantly synonymous substitutions (**Table 3**). The Mcd region showed the highest level of variability, particularly in the Mcd/NScd double mutant. Mixed viral populations were detected in all CD viruses, with most sequence variation concentrated within the first 180 codons of the modified M region (**Sup Fig 1**). Two recurrent amino acid substitutions were identified in Mcd-containing viruses: V1027I in the Gc protein and A613V in the Gn protein cytoplasmic tail. Both substitutions were conservative in nature but affected highly conserved residues. No amino acid changes were detected in rWT or NScd viruses. Overall, the sequencing results indicate partial genetic adaptation of codon-deoptimized viruses while maintaining the engineered attenuation phenotype.

**Table 3.** Summary of genome coverage and sequencing depth across viral segments.

| VIRUS | CLONE | SEGMENT | Genome Coverage (%) | Mean Coverage (X) | Length (nt) |
| --- | --- | --- | --- | --- | --- |
| **CDM** | **1** | **L** | 100,0% | 20,42 | 6404 |
|  |  | **M** | 100% | 5410,78 | 3885 |
|  |  | **S** | 95,0% | 20,55 | 1691 |
|  | **2** | **L** | 99,9% | 18,89 | 6404 |
|  |  | **M** | 100% | 4250,88 | 3885 |
|  |  | **S** | 96,6% | 18,84 | 1691 |
| **CDS** | **1** | **L** | 100% | 16,16 | 6404 |
|  |  | **M** | 100% | 38,13 | 3885 |
|  |  | **S** | 77,4% | 7,25 | 1691 |
|  | **2** | **L** | 97,3% | 6,76 | 6404 |
|  |  | **M** | 98,6% | 19,43 | 3885 |
|  |  | **S** | 99,1% | 4,59 | 1691 |
| **CDM/S** | **1** | **L** | 100% | 52,73 | 6404 |
|  |  | **M** | 100% | 14669,17 | 3885 |
|  |  | **S** | 98% | 3737,77 | 1691 |
|  | **2** | **L** | 100% | 31,93 | 6404 |
|  |  | **M** | 100% | 13569,30 | 3885 |
|  |  | **S** | 100% | 2718,14 | 1691 |
| **WT** | **1** | **L** | 100% | 262,28 | 6404 |
|  |  | **M** | 100% | 189,08 | 3885 |
|  |  | **S** | 100% | 71,97 | 1691 |

## DISCUSSION

Recently, the use of suboptimal codon pair bias has emerged as a novel strategy for generating attenuated viruses that can be used as LAV candidates. This approach has been successfully applied to poliovirus [39,40], influenza A virus [17,41,42], and respiratory syncytial virus [43], among others. However, codon pair deoptimization requires computational algorithms to redesign viral genomes, and different combinations of deoptimized codon pairs result in varying degrees of attenuation [39–43]. Thus, each codon pair-deoptimized virus must be empirically evaluated. To overcome these limitations, we explored an alternative strategy in which each amino acid residue of a viral protein is encoded by the least preferred codon in mammalian cells. Here, we provide the first evidence supporting the feasibility of this approach for attenuation of a bunyavirus RVFV. Recombinant RVFV encoding CD versions of the M or NSs genes replicated to high titers but exhibited reduced fitness *in vitro* and/or *in vivo*. Compared to rWT, replication of rMcd and rMcd/NScd was impaired in mammalian cells and, to a lesser extent, in insect cells, whereas rNScd replication was largely unaffected. *In vivo* studies revealed that rNScd and rMcd/NScd were highly attenuated in immunocompetent mice, while rMcd displayed moderate attenuation, indicating that CD of NSs has a stronger effect on virulence, with additional contribution from M gene deoptimization.

The attenuated phenotype observed with rNScd could be the consequence of the reduced ability of this virus to suppress the host innate antiviral response [35,44,45]. This hypothesis was assessed in a functional assay in vitro and using IFNAR^−/−^ mice. Interestingly, all CD viruses exhibited similar virulence in IFNAR^−/−^ mice, suggesting that attenuation is linked to impaired antagonism of innate immune responses. The underlying mechanisms remain to be elucidated and may involve reduced translation efficiency, altered RNA secondary structures, or those derived from changes in the increased CpG/UpA dinucleotide frequencies, as reported for other CD viruses [46].

The lack of attenuation in IFNAR^−/−^ mice underscores the need to explore additional strategies (if a higher level of attenuation is desirable), including CD of other viral genes. Importantly, CD-based attenuation can be combined with other previously described approaches that result in viral attenuation, such as targeted mutations in viral segments, to generate safer LAV [27,36]. If compared to conventional LAV approaches that rely on a few amino acid substitutions, the CD strategy described here offers unique advantages including, among others, the unlikely reversion of a CD virus to a WT virulent phenotype, based on the elevated number of nucleotide changes introduced in the CD approach or, the preservation of the full antigenic repertoire of the CD virus. In addition, NGS findings suggest that codon deoptimization of segment M promotes the emergence of adaptive mutations, with recurrent conservative substitutions in functionally relevant regions, potentially reflecting selective pressures to maintain essential protein structure and viral fitness. Overall, these data suggest that codon deoptimization of segment M has a greater impact on viral fitness, leading to increased genetic variability. Furthermore, these findings support the idea that codon deoptimization can be used as a tool to identify genomic regions with limited tolerance for variation, potentially highlighting structurally or functionally critical domains. However, this hypothesis requires further experimental validation.

## AUTHORS CONTRIBUTIONS

Conceptualization: A.N., L.M-S., and A.B.; Methodology: S.M., A.C., C.A., G.L., C.C., B.B., and A.N.; Data collection and interpretation: S.M., A.C., C.A., and A.N.; Funding acquisition and resources: A.N., L.M-S., B.B and A.B.; Writing, review, and editing: all authors have read and agreed to the published version of the manuscript.

## ACKNOWLEDGMENTS/FUNDING STATEMENT

This study was partially supported by the European Union’s Horizon Europe Research and Innovation Programme (grant agreement 101046133) and by grants PID2023-146428OB-I00, PID2024-158513OB-I00 and PID2021-122567OB-I00 funded by MCIN/ AEI/10.13039/501100011033/and by ERDF “A way of making Europe.” C.A. is enrolled on the PhD program of Microbiology at the Universidad Autónoma de Madrid.

